# Overabundance of *Asaia* and *Serratia* bacteria is associated with deltamethrin insecticide susceptibility in *Anopheles coluzzii* from Agboville, Côte d’Ivoire

**DOI:** 10.1101/2021.03.26.437219

**Authors:** Bethanie Pelloquin, Mojca Kristan, Constant Edi, Anne Meiwald, Emma Clark, Claire L. Jeffries, Thomas Walker, Nsa Dada, Louisa A. Messenger

## Abstract

**Background:** Insecticide resistance among mosquito species is now a pervasive phenomenon, which threatens to jeopardise global malaria vector control efforts. Evidence of links between the mosquito microbiota and insecticide resistance is emerging, with significant enrichment of insecticide degrading bacteria and enzymes in resistant populations. Using 16S rRNA amplicon sequencing, we characterised and compared the microbiota of *Anopheles* (*An.*) *coluzzii* in relation to their deltamethrin resistance and exposure profiles.

**Results:** Comparisons between 2-3 day old deltamethrin resistant and susceptible mosquitoes, demonstrated significant differences in microbiota diversity (PERMANOVA, pseudo-F = 19.44, p=0.0015). *Ochrobactrum, Lysinibacillus* and *Stenotrophomonas* genera, each of which comprised insecticide degrading species, were significantly enriched in resistant mosquitoes. Susceptible mosquitoes had a significant reduction in alpha diversity compared to resistant individuals (Shannon index: H=13.91, q=0.0003, Faith’s phylogenetic diversity: H=6.68, q=0.01), with *Asaia* and *Serratia* dominating microbial profiles. There was no significant difference in deltamethrin exposed and unexposed 5-6 day old individuals, suggesting that insecticide exposure had minimal impact on microbial composition. *Serratia* and *Asaia* were also dominant in 5-6 day old mosquitoes, regardless of exposure or phenotype, and had reduced microbial diversity compared with 2-3 day old mosquitoes.

**Conclusions:** Our findings revealed significant alterations of *An. coluzzii* microbiota associated with deltamethrin resistance, highlighting the potential for identification of novel microbial markers for insecticide resistance surveillance. qPCR detection of *Serratia* and *Asaia* was consistent with 16S rRNA sequencing, suggesting that population level field screening of the bacterial microbiota may be feasibly integrated into wider resistance monitoring if reliable and reproducible markers associated with phenotype can be identified.

## Background

Malaria remains a considerable public health problem with an estimated 229 million cases worldwide, including 409,000 deaths in 2019 alone[1]. Malaria mortality has fallen since 2010, largely due to the scale-up of treatment, diagnostics and insecticide-based vector control interventions, principally long-lasting insecticidal nets (LLINs) and indoor residual spraying (IRS). However, global gains in malaria control have begun to stall[2]. Insecticide resistance among major malaria vector species is now a pervasive phenomenon, affecting more than 90% of countries with ongoing transmission[2]. Of particular concern is the continued spread of resistance to pyrethroids, which were until recently, the only class of insecticide recommended for use in LLINs. Pyrethroids are still a crucial component of next-generation LLINs[3], and resistance may severely threaten the long-term effectiveness of contemporary vector control programmes.

Control of insecticide-resistant vector populations is predicated on a clear understanding of the complex interplay between molecular mechanisms and fitness costs which contribute to mosquito behaviour, phenotype and vectorial capacity, the genetic and local environmental factors driving ongoing resistance selection and the implications of resistance for intervention operational efficacy. Substantial progress has been made elucidating key target site mutations[4]–[7], over-expression of detoxification enzymes[8]–[12] and alternate gene families and pathways[13]–[17], all of which play important roles in resistance modulation. Furthermore, the recent publication of genome data for more than 1000 *Anopheles* (*An*.) *gambiae* sensu lato (s.l.) has illustrated the considerable genetic diversity among natural vector populations, raising concerns for the rapid evolution and spread of novel resistance mechanisms[18], [19].

In addition to host-mediated resistance mechanisms, evidence is emerging that changes in mosquito microbiota may confer resistance to certain insecticides. The mosquito microbiota is a heterogenous and variable network of microorganisms, comprising the bacterial, archaeal, viral, fungal, and other eukaryotic microbial communities which inhabit the mosquito cuticle and internal structures such as the midgut, salivary glands, and ovaries. Constituents of the microbiota can be either inherited from mother to offspring[20] or acquired from the environment, predominantly the larval habitat[21]. Characterisation of the microbiota in mosquitoes has shown varied phenotypic impacts on the host species including on fitness[22], blood feeding[23], fecundity[24], immunity[25], [26], pathogen infection[27]–[33] and transmission[34]. There is increasing interest in investigating symbionts of mosquito vectors because they may offer unique transmission-blocking opportunities. Similarly, studies on the role played by mosquito symbionts in insecticide resistance may offer a better understanding of the underlying mechanisms, and the potential for designing innovative control techniques[35] and developing new insecticide resistance monitoring tools.

The interaction between insecticide resistance and arthropod microbiota has been examined principally in agricultural pest species. Chlorpyrifos and fipronil resistant strains of the Diamondback moth, *Plutella xylostella* were shown to have a higher proportion of *Lactobacillales, Pseudomonadales* and *Xanthomonadales* bacteria[36]. Furthermore, the bean bug *R. pedertris* and allied stinkbug species harbour symbiotic *Burkholderia* bacteria which degrade fenitrothion, and are present in greater abundance when this insecticide is applied to their habitat.[37] As advanced molecular technologies become increasingly accessible, research in this area is expanding to disease vectors, with recent studies on several mosquito species. Whole metagenome sequencing of microbiota from wild-caught fenitrothion resistant and susceptible *An*. *albimanus* mosquitoes showed distinct differences between these two groups^65^. Fenitrothion-resistant mosquitoes had significant enrichment of organophosphate degrading bacteria and enzymes such as hydrolases, carboxylesterases and phosphomonoesterases. Resistant mosquitoes also had lower bacterial diversity, with an overabundance of *Klebsiella spp.* and a reduction in the relative abundance of *Enterobacter spp.* It was suggested that selection for organophosphate degrading bacteria may have developed alongside resistance, potentially in response to prior insecticide exposure[38]. F_1_ progeny of field-caught *An. albimanus* exposed to the pyrethroids alpha-cypermethrin and permethrin had significantly greater abundance of bacteria from the genus *Pseudomonas*, of which several strains have been shown to metabolise pyrethroids, and from the genus *Pantoea*[39], which had previously been identified in insecticide-resistant mosquitoes[38]. *Pseudomonas*, alongside *Clostridium* and *Rhizobium* species, were also implicated in lambda-cyhalothrin resistance in wild populations of *Aedes (Ae.) aegypti* from Colombia[40]. Addition of tetracycline to temephos-resistant strains of *An. stephensi* destroyed the bacterial component of the microbiota and significantly reduced the activity of three main resistance enzymes: α esterase, glutathione-S-transferase, and acetylcholinesterase, restoring mosquito susceptibility[41]. Similarly, sterilisation of *An. arabiensis* gut microbiota by antibiotics resulted in a decreased tolerance to deltamethrin and malathion[42].

To date, information on field populations of the *An. gambiae* complex, the main malaria vectors in sub-Saharan Africa, is limited to recent reports of significant enrichment of known pyrethroid degrading taxa (*Sphingobacterium, Lysinibacillus* and *Streptococcus*) in permethrin-resistant *An. gambiae* sensu stricto (s.s.) from Kenya[43]. To address this deficit, we comparatively characterised the bacterial microbiota of *An. coluzzii,* collected from an area of high pyrethroid resistance in Côte d’Ivoire. We specifically focused on determining the effects of deltamethrin resistance intensity on host microbiota and identifying any microbial taxa associated with resistance phenotypes.

## Methods

### Mosquito collections and mass rearing

This study was conducted in Agboville (GPS: 5°55’21” N 4°13’13” W), Agnéby-Tiassa region, south-east Côte d’Ivoire. The location was chosen because of its high mosquito densities, malaria prevalence (26% in children <5 years in recent estimates[44]) and intense deltamethrin resistance[45]. The main industry is agriculture, with livestock such as cows, goats and chickens living close to households and cultivation of crops including bananas, cocoa and rice[46].

Sampling was conducted between 5th July and 26th July 2019, coinciding with the long rainy season (May-November) and peak malaria transmission. Adult mosquitoes were collected using HLCs, inside and outside households from 18:00h to 06:00h. Fieldworkers used individual haemolysis tubes to collect host-seeking mosquitoes, which were transported each morning to the Centre Suisse de Recherche Scientifique en Côte d’Ivoire (CSRS) in Abidjan. Blood-fed mosquitoes, morphologically identified as female *An. gambiae* s.l.[47], were transferred to cages with 10% sugar solution and left for 2-3 days to become fully gravid.

Five hundred and eighty fully gravid females were used for forced oviposition. Oviposition was achieved by placing a single gravid mosquito into an 1.5 ml Eppendorf tube, half filled with damp cotton wool, with small holes in the tube cap for ventilation[48]. Mosquitoes were held under standard insectary conditions (25°C, 70% humidity and a 12-hour light-dark cycle) until eggs were laid or adult death. Eggs were removed daily and placed into sterile paper cups containing distilled water and NISHIKOI (Nishikoi, United Kingdom)[49] staple fish food pellets. Emergent larvae were reared in 50 cm washing up bowls, in distilled water under the same insectary conditions. Pupae were removed daily and separated by sex with the aid of a stereomicroscope. Female pupae were put in a clean plastic cup with distilled water and placed in a cage for eclosion, while male pupae were discarded. Adults were housed in cages in an incubator (26.6°C, 70% humidity) with a 12-hour light-dark cycle and given unlimited access to 10% glucose solution. The cages were checked to ensure that only virgin females were used in bioassays, as mating can potentially introduce changes to the microbiome[20]. Care was also taken to ensure that no mosquito obtained a blood meal during handling, as this can significantly decrease bacterial diversity in the gut[50].

### Determining deltamethrin resistance status of adult F1 progeny of field-caught An. gambiae s.l

Deltamethrin resistance was characterised using Centre for Disease Control (CDC) bottle bioassays[51], with some modifications. Two to three-day old (d) virgin F_1_ females were exposed to 1, 5 or 10 times the diagnostic dose of deltamethrin (12.5μg/bottle) for 30 minutes. Stock solutions of deltamethrin were prepared using 100% ethanol as the solvent. Per bioassay, multiple 250mL Wheaton bottles, and their lids, were coated with 1mL stock solution and left to dry in a dark storage area to avoid exposure to UV light. A control bottle, treated with 1mL ethanol, was assayed in parallel. Prior to bioassay testing, approximately 20-25 mosquitoes were aspirated into holding cups. After 1-2 hours of acclimatisation, they were introduced into each test or control bottle.

Knock-down was scored at 0, 15 and 30 minutes. A subset of mosquitoes which were alive at 60 minutes were held for 72 hours, with mortality recorded every 24 hours. These were housed in paper cups in the insectary, with unlimited access to sterile 10% glucose made with distilled water. Mosquitoes were counted as dead if they were unable to stand as per WHO criteria[51].

At the end of the bioassay and subsequent holding time, mosquitoes were classified as: susceptible if they were knocked-down following exposure to 1x deltamethrin, resistant if they survived 60 minutes or 72 hours post-exposure to 1x, 5x or 10x deltamethrin, or controls if they were in the ethanol coated bottle. Specimens were separated into their respective phenotype and concentration/time group and stored at −70°C.

### DNA extraction

DNA was extracted from 380 mosquitoes which had been categorised as resistant, susceptible or unexposed to deltamethrin. Individuals were homogenised in a QIAGEN® TissueLyser II with sterilised 5mm stainless steel beads for 5 minutes at 30hz/sec and incubated overnight at 56°C. DNA was extracted using a QIAGEN DNeasy® 96 Blood and Tissue Kit (Qiagen®, UK) as per the manufacturers protocol[52] with DNA eluted in 45μL of buffer AE. Extracted DNA was stored at −70°C.

Four blank extraction controls were processed alongside mosquitoes: three blanks containing RNase-free water as the extraction template and one blank containing the 70% ethanol used for reagent dilution and sterilisation of instruments. All steps were performed under sterile conditions, with tweezers and other instruments being rinsed with 70% ethanol in between handling each mosquito, to avoid microbial or DNA contamination.

### PCR for mosquito species identification

Individual mosquitoes were identified to species level according to Santolamazza et al[53]. PCR reactions contained 2μL of 10μM forward primer (5’-TCGCCTTAGACCTTGCGTTA-3’), 2 μL of 10μM reverse primer (5’-CGCTTCAAGAATTCGAGATAC-3’), 1μL extracted DNA and 10μL HotStart Taq Master Mix (New England Biolabs, UK), for a final reaction volume of 20μL. Prepared reactions were run on a BioRad T100™ thermal cycler with the following conditions: 10 minutes denaturation time at 94°C, followed by 35 amplification cycles of 94°C for 30 seconds, 54°C for 30 seconds and 72°C for 60 seconds, followed by a final extension at 72°C for 10 minutes. PCR products were visualised on 2% E-gel agarose gels in an Invitrogen E-gel iBase Real-Time Transilluminator. A Quick-Load^®^ 100bp DNA ladder (New England Biolabs, UK) was used to determine band size. Amplified PCR products of 479 bp or 249 bp were indicative of *An. coluzzii* or *An. gambiae* s.s., respectively. As the dominant species, only *An. coluzzii* individuals of the same age and resistance phenotype were selected and pooled for 16S rRNA sequencing.

### 16S rRNA gene amplicon sequencing

DNA concentration from each mosquito was measured using an Invitrogen Qubit™ 4 Fluorometer (Thermo Fisher Scientific, USA). Pools were prepared by combining equal concentrations of DNA from 3 mosquitoes of the same phenotype/deltamethrin concentration/time group to give 100ng in a final volume of 20μL (Table S1). Two negative controls, one comprised of a pool of the three RNase-free water blanks mentioned above, and the 70% ethanol blank, were processed in parallel.

The microbial composition of the microbiome was determined by amplification of the V3-V4 region of the *16S rRNA* gene, using the following primers: 5’-CCTACGGGNGGCWGCAG-3’ and 5’-GGACTACHVGGGTATCTAATCC-3’. PCR reactions were prepared in a 25μL reaction volume, comprising 12.5μL of KAPA^©^ HiFi Hot Start ReadyMixPCR Kit[54] (Roche, Switzerland), 0.5μL of forward and reverse primers (10μM) and 12.5ng DNA. The following PCR cycling was used: 95°C for 3 minutes, 35 cycles of 95°C for 30 seconds, 55°C for 30 seconds and 72°C for 30 seconds, followed by a final extension at 72°C for 5 minutes. The resulting amplified PCR products were purified with AMPure XP beads (Beckman Coulter, UK) at 1x sample volume.

Next an index PCR was performed using 5μL purified PCR products, 5μL of Nextera XT Index 1 Primers (N7XX), 5μL of Nextera XT Index 2 Primers (S5XX) (both from the Nextera XT Index kit, Illumina, USA), 10μL PCR grade water and 25 μL of KAPA^©^ HiFi Hot Start ReadyMix (Roche, Switzerland). The following PCR cycling was used: 95°C for 3 minutes, 12 cycles of 95°C for 30 seconds, 55°C for 30 seconds and 72°C for 30 seconds, followed by a final extension at 72°C for 5 minutes. The final library was purified with AMPure XP beads, at 1.12x sample volume, before quantification.

Sequencing was performed on an Illumina^©^ MiSeq^©^ platform. Libraries were sequenced as 250 bp paired-end reads. Sequences were demultiplexed and filtered for read quality using Bcl2Fastq conversion software (Illumina, Inc.). In total, 1,156,076 sequences were generated in the FASTQPhred33 format44.

### Data cleaning and filtering

Sequencing data were imported into the ‘Quantitative Insights Into Microbial Ecology’ pipeline, version 2020.8 (Qiime2)[55], and primary analysis was performed on the reverse reads, as the quality of the forward reads were not sufficient for merging (Figure S1). Sequencing primers and adapters were removed using the ‘cutadapt’ plugin[56] with an error rate of 10%. The divisive amplicon denoising algorithm (DADA2) plugin[57] was used to ‘denoise’ sequencing reads, removing phiX reads and chimeric sequences, to produce high resolution, ASVs[58]. DADA2 was run using the denoise-single command, with samples truncated at 206 nucleotides (trunc-len 206), to remove bases with a low-quality score. All other parameters were set to default. The resulting feature table[59] and sequences were filtered to remove ASVs present in the two blank samples and those with a frequency of below 100 to reduce biases in comparison of diversity indices across groups, and especially in differential abundance tests.

### Taxonomic annotation

Taxonomic annotation of ASVs was performed using the -feature-classifier plugin[60], with a Naïve-Bayes classifier[61] pretrained on the *16S* SILVA reference (99% identity) database version 132. The -extract-reads command was used to trim the reference sequences to span the V3-V4 region (425bp) of the *16S rRNA* gene. Any features not classified to phylum level were also removed, these included hosts’ mitochondrial *16S rRNA* genes. The resulting ASV table was exported into R (version 3.6.3) for analysis with the phyloseq package.

### Bacterial diversity analysis

A rooted and unrooted phylogenetic tree was generated using the qiime phylogeny plugin[62],[63],[64] and were used to compute alpha and beta diversity metrics using the qiime2-diversity[65] plugin. For alpha diversity metrics, samples were rarefied[66] at a depth of 2359; where alpha rarefaction curves plateaued, indicating that there was adequate sampling of the microbiota during sequencing. Beta diversity metrics were computed for both rarefied and non-rarefied data, with no significant differences between methods (Table S2); non-rarefied data are presented herein. 2-3 day old and 5-6 day old mosquitoes were analysed separately, as age was shown to significantly impact the bacterial composition of the microbiota.

Two methods of alpha diversity were selected: Shannon diversity Index, which considers the abundance and evenness of ASVs present, and Faith’s Phylogenetic Diversity, a measure of community richness which incorporates phylogenetic relationships between species. Pairwise Kruskal-Wallis comparisons of these alpha diversity indices between groups of insecticide resistance phenotypes were performed, with Benjamini-Hochberg false discovery rate (FDR) correction for multiple comparisons[67]. Significance was set to FDR adjusted *p* value i.e. *q* value < 0.05.

Bray-Curtis Dissimilarity Index[68][69], which measures differences in relative species composition between samples, was chosen as the beta diversity metric. Comparisons of this index between insecticide resistance phenotype groups were conducted using pairwise PERMANOVA tests with 999 permutations[70]. Results were visualised using PCoA generated using the phyloseq[71] package. Significance was set to *p* value < 0.05.

### Determination of association between microbiota composition and insecticide resistance phenotype, and identification of differentially abundant microbial taxa

Comparison of alpha and beta diversity indices indicated that both insecticide resistance phenotype and mosquito age affected the bacterial composition of *An. coluzzii* in this study. Following taxonomic annotation of ASVs, multinomial regression and differential abundance analysis was performed using Songbird[72] to determine the microbial taxa which were associated with and differentially abundant across insecticide resistance phenotype for mosquitoes separated by age group. Songbird is a compositionally aware differential abundance method which ranks features based on their log fold change with respect to covariates of interest[72] The following Songbird parameters were used: epochs 10000, number of random test examples 15, differential prior 0.5. The fit of the model was tested against the null hypothesis (-p-formula “1”). Differential log ratios of features were computed in Qurro[73]. We present the highest and lowest 10% ranked features associated with resistance phenotype. Analysis of Composition of Microbiome method (ANCOM) was used to complement Songbird analysis, and this was computed using the composition plugin[74] with all parameters set to default. Significance was determined using the automatic cut off for the test statistic, W[75].

### Quantitative PCR (qPCR) validation of sequencing data

The abundance of *Serratia spp.* and *Asaia spp.* was assessed using qPCR, relative to the nuclear single-copy *An. gambiae* s.l. ribosomal protein S7 housekeeping gene (*RPS7*). *Serratia* reactions contained 1μL of 10μM forward primer (5’-CCGCGAAGGCAAAGTGCACGAACA-3’), 1μL of 10μM reverse primer (5’-CTTGGCCAGAAGCGCACCATAG-3’)[76], 2μL of pooled DNA and 5μL LightCycler® 480 SYBR Green Master Mix (Roche, UK), for a final reaction volume of 10μL. Prepared reactions were run on an Agilent Technologies Stratagene Mx3005P qPCR system which performed 40 cycles of 95°C for 15 seconds and 60°C for 1 minute, followed by a dissociation curve. *Asaia* reactions contained 1μL of 10μM forward primer (5’-GCGCGTAGGCGGTTTACAC-3’), 1μL of 10μM reverse primer (5’-AGCGTCAGTAATGAGCCAGGTT-3’)[77], 2μL of pooled DNA and 5μL LightCycler® 480 SYBR Green Master Mix (Roche, UK), for a final reaction volume of 10μL. Prepared reactions were run on an Agilent Technologies Stratagene Mx3005P qPCR system with the following conditions: 95°C for 15 minutes, 40 cycles of 95°C for 10 seconds, 60°C for 10 seconds and 72°C for 10 seconds, followed by a dissociation curve. *RSP7* reactions contained 1μL of 10μM forward primer (5’-TCCTGGAGCTGGAGATGA AC-3’), 1μL of 10μM reverse primer (5’-GACGGGTCTGTACCTTCT GG-3’), [78]2μL of pooled DNA and 5μL LightCycler® 480 SYBR Green Master Mix (Roche, UK), for a final reaction volume of 10μL. Prepared reactions were run on an Agilent Technologies Stratagene Mx3005P qPCR system with the following conditions: 40 cycles of: 95°C for 10 seconds, 65°C for 60 seconds and 97°C for 1 second, followed by a dissociation curve. All samples were run in technical triplicate. Relative bacterial abundance was normalised relative to the endogenous control gene (*RPS7*). qPCR results were analysed using the MxPro software (Agilent Technologies).

## Results

### Species identification and deltamethrin resistance profiles

In total, 580 *An. gambiae* s.l. were collected from Agboville using human-landing catches (HLCs), during the rainy season in July 2019. Of these, 245 (42%) laid eggs via forced oviposition. Following larval development, 1015 F_1_ *An. gambiae* s.l. pupae were identified as female and tested in deltamethrin resistance intensity assays as 2-3 day old adults. Individuals were classified as susceptible if they were knocked-down following exposure to 1x deltamethrin, resistant if they survived 60 minutes (2-3 day old) or 72 hours (5-6 day old) post-exposure to 1x, 5x or 10x deltamethrin, or controls if they were unexposed to insecticide (comprising a mix of age-matched individuals of unknown phenotype). A total of 380 mosquitoes were randomly selected for DNA extraction, across all exposure and time groups, with 338 individuals identified as *An. coluzzii* (78.3%). From the remaining individuals, 31 were *An. gambiae* s.s. (8.1%), 10 failed to amplify (2.6%), and one individual was an *An. gambiae* s.s.–*An. coluzzii* hybrid (0.26%). Table S1 summarises the number of mosquitoes selected for DNA extraction, pooling and sequencing.

### Sequencing metrics

A total of 1,156,076 reverse reads were obtained from sequencing. Quality control and denoising resulted in 2,999 unique amplicon sequence variants (ASVs), 878,155 in total. Filtering of ASVs associated with water and ethanol blanks, low frequency ASVs and ASVs not classified to phylum level resulted in 210 unique ASVs, totalling 556,254 across 94 pools of mosquitoes. Table S3 summarises the number of sequences processed per sample and the number of reads remaining after denoising and filtering.

### Susceptible An. coluzzii had microbiota which were significantly different to, and less diverse than, resistant mosquitoes

Comparison of the Bray-Curtis dissimilarity index using pair-wise PERMANOVA with 999 permutations showed significant differences in bacterial composition between microbiota of 2-3 day old deltamethrin resistant and susceptible *An. coluzzii* (pseudo-F = 19.44, p=0.0015). Principal Coordinate Analysis (PCoA) visualisations showed the microbiota of susceptible mosquitoes clustered away from resistant and control mosquitoes (Figure 1), indicating that the microbiota of susceptible mosquitoes were more similar to each other than to resistant and control mosquitoes.

**Figure 1.**
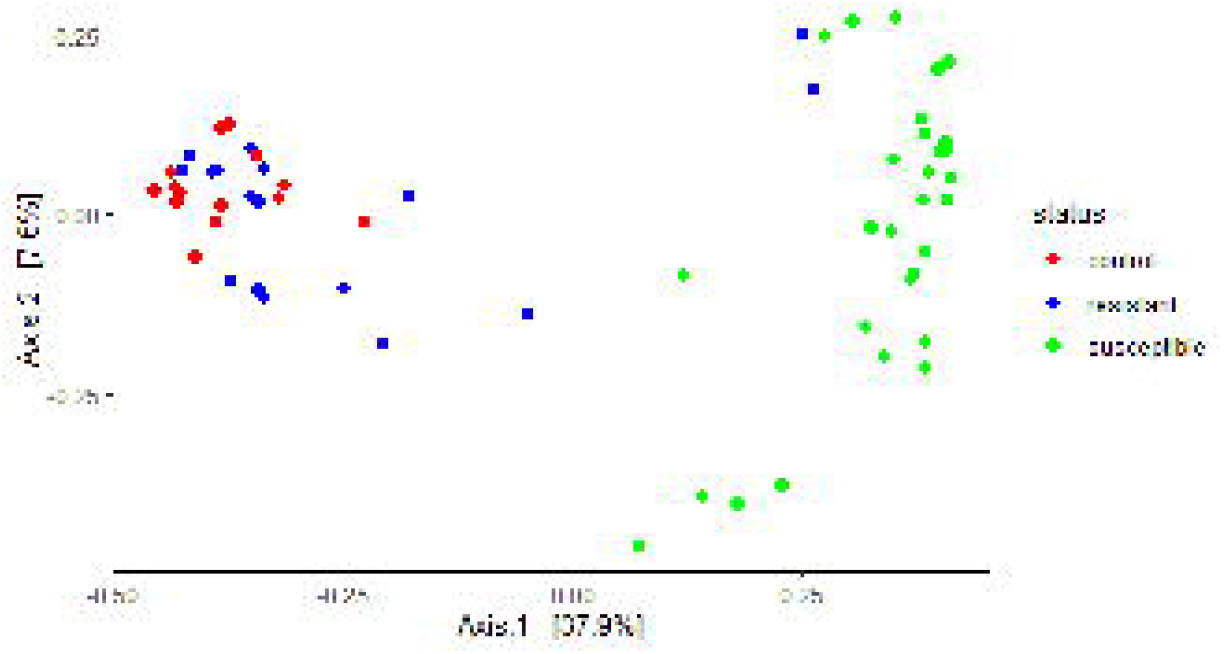
PCoA plot showing Bray Curtis distance of microbiota between resistant, susceptible and control F_1_ 2-3 day old adult *An. coluzzii*. Each point represents the bacterial composition of a pool of three mosquitoes of the same resistance phenotype. There was a distinct separation between resistant/control and susceptible mosquitoes, which was shown to be a significant difference using a pairwise PERMANOVA (999 permutations) (pseudo-F = 19.44, p=0.0015).

Susceptible mosquitoes had significantly lower Shannon and Faith Phylogenetic Diversity indices than resistant (Shannon: H=13.91, q=0.0003, Faith: H=6.68, q=0.01) and control mosquitoes (Shannon: H=22.6 q=0.000006, Faith: H = 16.6, q = 0.0001) of the same age, indicating that the susceptible group had reduced microbial diversity. There was no significant difference in alpha or beta diversity in deltamethrin exposed and unexposed 5-6 day old mosquitoes (Shannon: H=5.12, q=0.02, Faith: H=0.27, q=0.6, Bray-Curtis: pseudo-F=1.61, q=0.17), suggesting that insecticide exposure during the CDC bottle bioassays had minimal impact on microbial composition.

### Serratia and Asaia dominated in older, and younger susceptible An. coluzzii

Following taxonomic annotation of ASVs to the genus or lowest possible taxonomic level, 114 and 57 bacterial taxa were detected in 2-3 day old and 5-6 day old mosquitoes, respectively. The less diverse 5-6 day old microbiota was predominantly comprised of ASVs assigned to the genera *Serratia* (75.5%) and *Asaia* (13.6%) (Figure S2). In 2-3 day old mosquitoes, microbial composition varied by resistance phenotype. Control mosquitoes had the highest number of taxa present (n=97), followed by resistant (n=90) and susceptible (n=66). 20 taxa were unique to control mosquitoes, and 15 to resistant mosquitoes. No taxa were unique to the susceptible group of mosquitoes, and 60 taxa were common to all groups (Figure 2).

**Figure 2.**
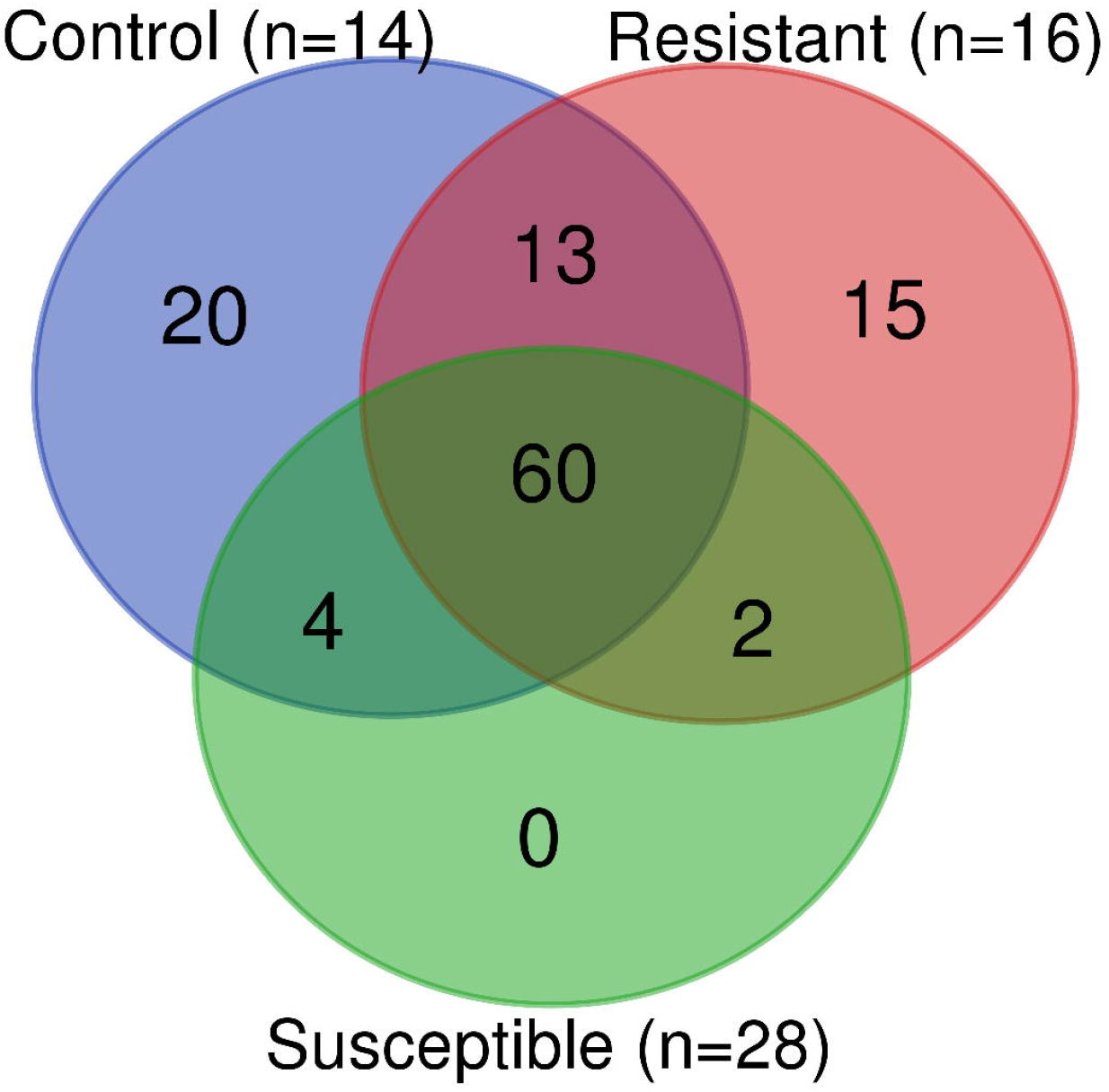
Venn diagram showing number of bacterial taxa unique to or shared between pools of 2-3 day old resistant, susceptible or control mosquitoes. Taxa were identified to genus level or lowest possible taxonomic rank. n=number of pools (each pool consists of three mosquitoes of the same age and phenotype).

In control mosquitoes, an unclassified species within the Enterobacteriaceae family (15.24%), *Acinetobacter* (8.83%) and *Staphylococcus* (8.29%) were most abundant, whilst Enterobacteriacea*e* (15.12%), *Acinetobacter* (14.26%) and *Serratia* (11.8%) were the most abundant in resistant mosquitoes. In susceptible mosquitoes, *Serratia* (56.4%) and *Asaia* (30.92%) were the dominant genera, with *Acinetobacter* (1.96%), Enterobacteriaceae (1.57%) and *Staphylococcus* (1.4%) present at low abundance (Figure 3). The remaining 61 taxa were present at an abundance of less than 1% of total ASVs present (Figure S2, Table S4).

**Figure 3.**
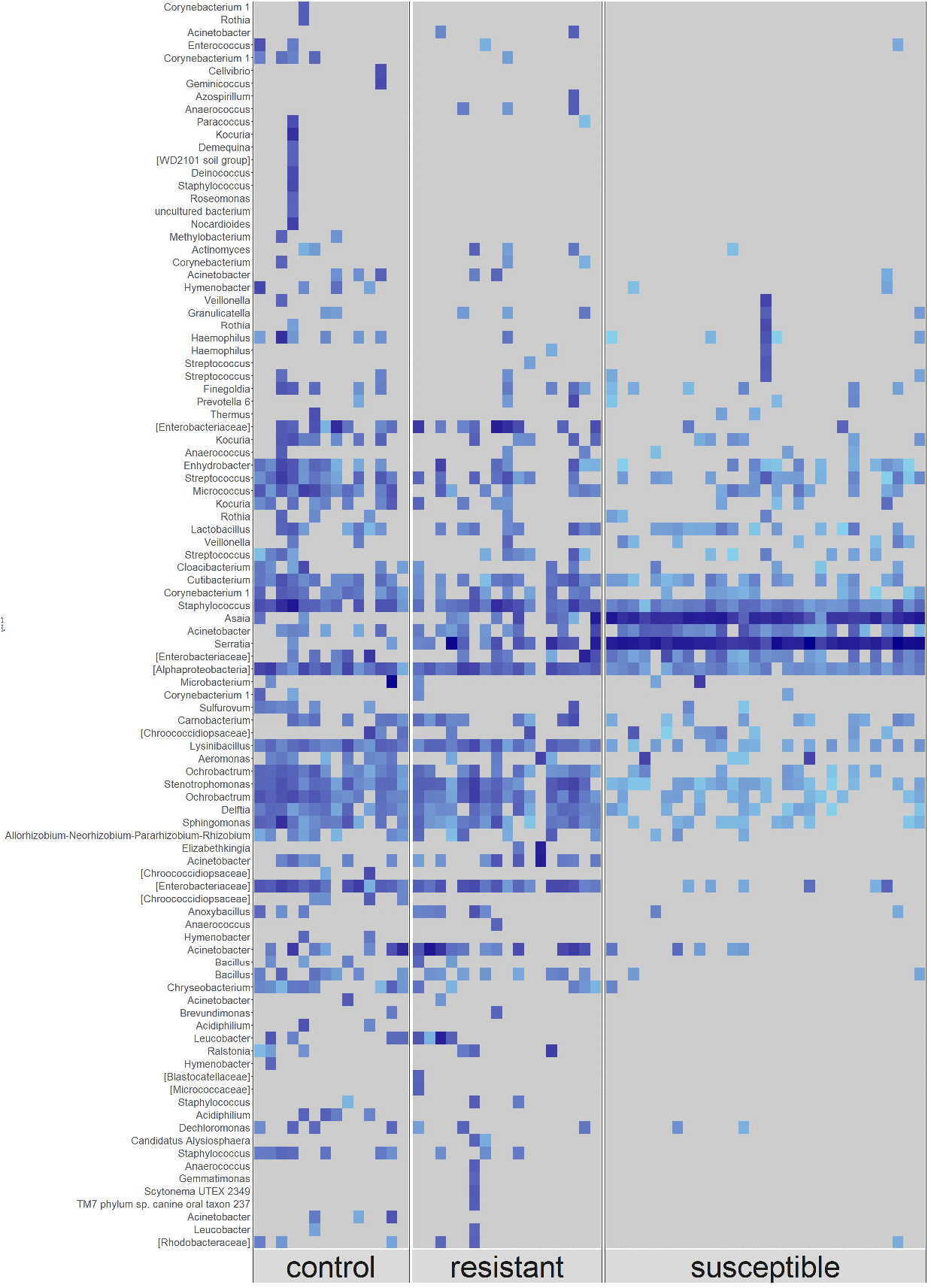
Raw frequency of ASVs from the microbiota of control (n=14), resistant (n=16) and susceptible (n=28) F_1_ 2-3 day old adult *An. coluzzii*. Each column represents a pool of three mosquitoes of the same phenotype. ASVs were annotated to genus level or lowest possible taxonomic level (in square brackets). Only taxonomically annotated ASVs with a frequency of >150 are shown. Light blue indicates a low frequency of ASVs present, whilst darker blue indicates a higher frequency. Grey indicates ASV not present in that pool.

### *Differential rankings confirmed Asaia and Serratia were significantly associated with susceptibility and Stenotrophomonas, Ochrobactrum, Lysinibacillus* and *Alphaproteobacteria were significantly associated with phenotypic resistance*

Songbird was used to identify taxa which were differentially abundant in 2-3 day old resistant, susceptible or control mosquitoes. Evaluation of our Songbird model with resistance phenotype as the variable, against a baseline model with no variable resulted in a pseudo Q-squared value of 0.42, indicating that the model had not been overfit and that roughly 42% of variation in the model was predicted by resistance phenotype (Figure S3). There were significant differences in the log ratios of highest to lowest ranked taxa between resistant and susceptible microbiota (Figure 4, Table S5), suggesting that the highest ranked taxa were significantly overabundant in resistant microbiota, and the lowest ranked taxa were significantly overabundant in susceptible microbiota.

**Figure 4.**
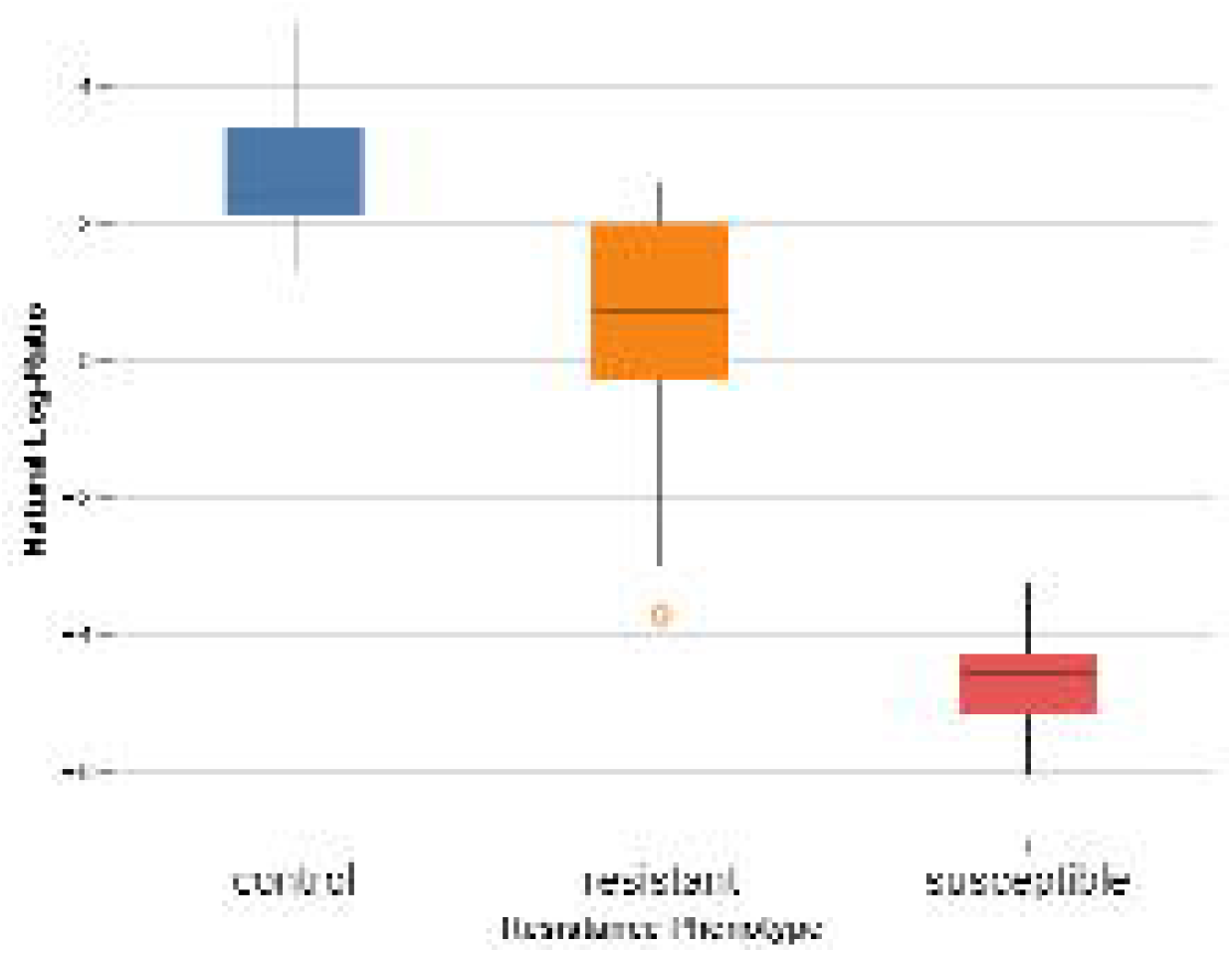
Log ratios of 10% highest ranked features to 10% lowest ranked features in control, resistant and susceptible 2-3 day old F_1_ *An. coluzzii*. Susceptible mosquitoes had a significantly lower ratio than control or resistant mosquitoes indicating that the lowest ranked features were overabundant in the susceptible group, whilst the highest ranked features were overabundant in either resistant or control mosquitoes.

*Stenotrophomonas, Ochrobactrum, Lysinibacillus* and *Alphaprotebacteria* (highest ranked) were most strongly associated with insecticide resistance whilst *Serratia, Aerococcus, E. shigella* and *Asaia* (lowest ranked) were most strongly associated with insecticide susceptibility (Figure 5). Comparing log ratios of control and susceptible pools indicated that *Rhodococcus, Sphingomonas, Haemophilus* and *E. shigella* were most strongly associated with controls, whilst an uncultured Chroocooccidiopsaceae, *Serratia,* an unclassified member of Enterobacteriacea and *Asaia* were most strongly associated with susceptible mosquitoes (Table S5).

**Figure 5.**
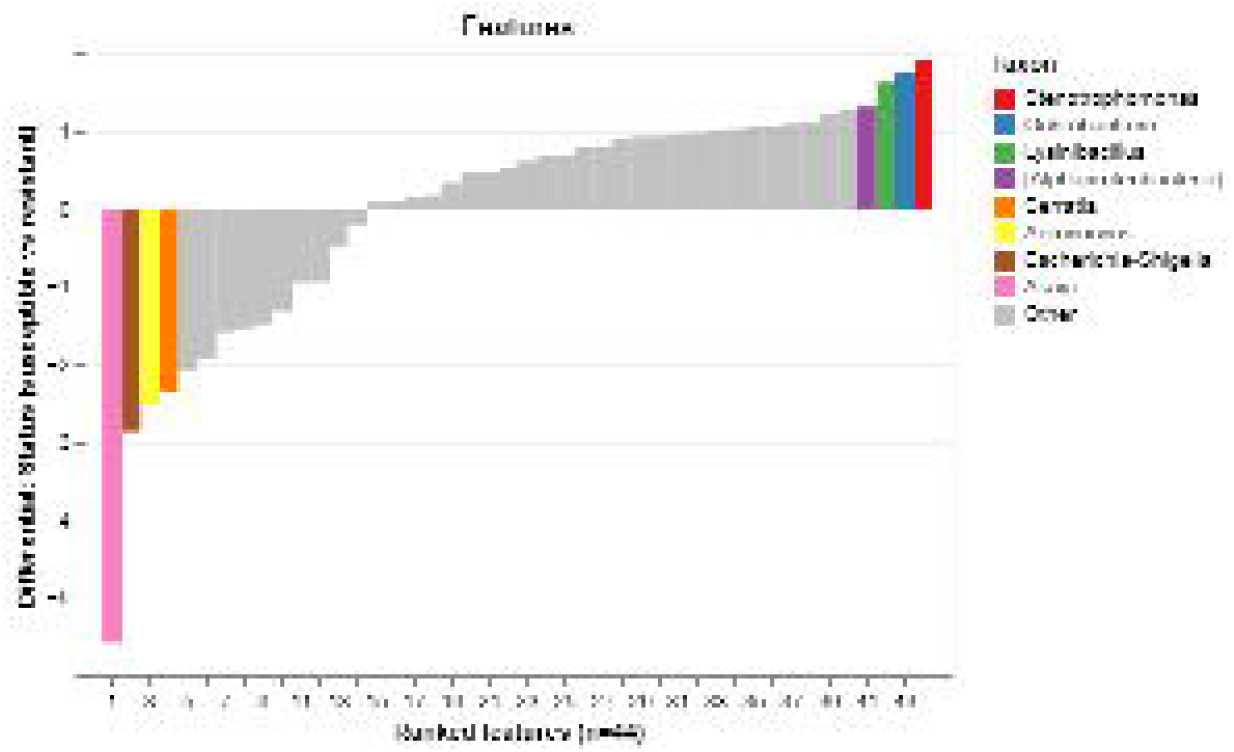
Sorted differential ranks of features associated with resistant or susceptible phenotype in 2-3 day old *An. coluzzii*. The highest 10% and lowest 10% of ranked features are shown, coloured by their corresponding assigned taxon. Taxa are shown to genus or lowest possible taxonomic level (square brackets).

These results were confirmed by the ANCOM method. *Serratia* (W=208) and *Asaia* (W=208) were significantly overabundant in susceptible mosquitoes relative to resistant and controls, whilst *Ochrobactrum* (W=199), *Lysinibacillus* (W=188) and *Enterobacteriaceae* (W=201) were overabundant in resistant and control mosquitoes (Figure S4).

### *Increased abundance of* Serratia *and* Asaia *species in susceptible individuals confirmed by qPCR*

Quantitative PCR assays confirmed that *Serratia* was significantly overabundant in 2-3 day old susceptible mosquitoes compared to deltamethrin resistant (5x p=0.028, 10x p=0.002) and control (p=0.02) (average CT value for susceptible: −8.4 [95% CI: −9.0 – −7.74]; 5x: −7.43 [−7.9 – −6.9]; 10x −6.3 [−7.2 – −5.4]; control: −7.2 [−7.9 – −6.6]). *Asaia* was also significantly overabundant in 2-3 day old susceptible mosquitoes compared to resistant (5x p= >0.001, 10x p<0.001) and control (p<0.001) (average CT values for susceptible: −7.6 [95% CI: −8.5 – −6.8]; 5x: 0.8 [−1.6 – 3.2]; 10x: 5.1 [3.0 – 7.1]; control: 4.5 [3.3 – 5.7]). Five to six day old mosquitoes also had increased abundance of both bacterial species (average CT value for 5-6 day old 5x: S*erratia* −8.2 [95% CI: −8.9 – −7.4] *Asaia* −1.3 [−5.6 – 2.9]; 5-6 day old 10x: *Serratia* −7.5 [−7.9 – −7.1] *Asaia* 5.4 [4.6 – 6.3]; 5-6 day old control *Serratia* −8.7 [−9.6 – −7.8] and *Asaia* −0.68 [−3.0 –1.7].

## Discussion

There is increasing evidence for an association between insecticide resistance phenotype[38], [43], [79] and *Anopheles spp.* microbiota. This study revealed distinct differences between the microbiota of deltamethrin resistant and susceptible *An. coluzzii*, with significant enrichment of insecticide degrading taxa in resistant individuals and an overabundance of *Serratia* and *Asaia* taxa in susceptible individuals. This population of field-caught *An. gambiae* s.l. from Agboville, South-East Côte d’Ivoire has previously been characterised as intensely resistant to pyrethroids, with average vector mortality of 14.56% [95% CI: 8.92-22.88%], 61.62% [95% CI: 51.58-70.75%] and 73.79% [95% CI: 64.35-81.45%] to one, five and ten times the diagnostic dose of deltamethrin, respectively, and pyrethroid resistance associated with over-expression of CYP450 enzymes (*CYP6P4*, *CYP6Z1* and *CYP6P3*)[45].

Our study demonstrated significant differences in alpha and beta diversity between deltamethrin-resistant and susceptible *An. coluzzii.* Resistant mosquitoes harboured a wider variety of microbial taxa and had more microbial diversity both within and between themselves. Susceptible mosquitoes had fewer bacterial taxa and were far more homogenous, with *Serratia* and *Asaia* dominating in all samples. Previous studies have demonstrated differences between the microbiota of insecticide resistant and susceptible *An. stephensi*[38]*, An. arabiensis*[42], and *An. gambiae* s.s[43]. This study is the first to detect an increase in alpha diversity in resistant mosquitoes with other studies reporting no difference[39], [43] or a decrease[80]. There are multiple potential reasons for the decreased microbial diversity previously identified, including processing of individuals rather than pooled mosquitoes, host species-specific differences, and variability among geographical collection sites[80]. Furthermore, field-caught mosquitoes may be of unknown age and physiological status; prior environmental insecticide exposure might also be responsible for reducing overall diversity, as bacteria with the ability to metabolise insecticides have greater access to compounds for growth and reproduction and can outcompete other species[38]. In our study, larvae were raised in distilled water, adults were age-standardised, unable to mate or blood-feed and had no insecticide exposure prior to resistance assays. We therefore consider this inherited microbiota to be linked to the resistance status of the host, with mosquitoes characterised by a wider range of bacteria having an increased chance of harbouring an insecticide degrading strain.

The role of bacteria in the degradation of pesticides has been widely studied[36], [37], [81], mainly due to the interest in bioremediation – use of bacteria to restore pesticide contaminated soils and water. However, emerging evidence that these bacteria could also contribute to pesticide resistance is concerning. Our study identified significant enrichment of several insecticide degrading taxa in resistant compared with susceptible mosquitoes: *Ochrobactrum, Lysinibacillus* and *Stenotrophomonas*. *Ochrobactrum spp.* have been isolated from contaminated soil, and shown to degrade a variety of insecticides including pyrethroids[82], [83] and organophosphates[84], [85]. Similarly, *Lysinibacillus spp.* derived from soil and sewage can metabolise deltamethrin[86] and cyfluthrin[87] whilst *Stenotrophomonas* in the microflora of cockroaches living in pesticide treated environments can degrade endosulfan *in vitro*[88]. Elevated expression of xenobiotic degrading genes[86] and enzymes[38] may be contributing to the insecticide degrading properties of these bacteria as well as direct degradation of pesticides.

While certain species of bacteria can confer insecticide resistance to the host, others may influence susceptibility. Indigenous gut bacteria have been implicated in the susceptibility of the gypsy moth, *L. dispar*, to the insecticidal toxin *Bacillus (B.) thuringiensis*. Treatment of larvae with antibiotics eliminated gut microbiota, and subsequently reduced mortality to *B. thuringiensis*; susceptibility was restored upon oral administration of *Enterobacter sp.*, a gram negative bacterium widely present in the *L. dispar* gut[89]. In our study, *Serratia* and *Asaia* were found to be significantly overabundant in susceptible mosquitoes. Whilst there are no prior reports of an association of these species with mosquito insecticide resistance phenotype, when the relative abundance of *Serratia sp.* in the gut of the diamondback moth was increased, susceptibility to chlorpyrifos significantly increased[81]. *Serratia marcescens* plays a role in the susceptibility of field-caught *Aedes* to dengue virus infection by secreting *Sm*Enhancin, an enzyme which digests gut epithelia mucins, enabling the virus to penetrate the gut[35]. The Bel protein, similar to *Sm*Enhancin, and produced by the bacteria *B. thuringiensis,* has been shown to significantly increase the toxicity of Cry1Ac toxin in the cotton bollworm larvae, *Helicoverpa armigera*; by perforating the midgut peritrophic matrix and degrading the insect intestinal mucin, enabling the toxin to reach the target epithelial membrane[90]. A similar mechanism may occur in these mosquitoes, whereby proteins produced by *Serratia spp.* increase the permeability of the internal organs to deltamethrin, enabling it to reach its target in the mosquito nervous system.

*Asaia* and *Serratia* sp. have also both previously been implicated in modulation of *Anopheles* vector competence. *Asaia sp.* have been shown to activate antimicrobial peptide expression in *An. stephensi*[91] while some strains of *S. marcescens* isolated from *An. sinensis* can inhibit *Plasmodium* development by altering the immunity-related effector genes TEP1 and FBN9[27] *Serratia* spp. may also directly inhibit malaria parasite development by secretion of serralysin proteins and prodigiosin, which can have a pathogen-killing effect *in vitro*[92]; the latter can also act as a larvicidal agent against *Ae. aegypti* and *An. stephensi*[93]. By comparison, the presence of a dominant commensal Enterobacteriaceae has been positively correlated with *Plasmodium* infection[33]. Elucidating the impact of mosquito host microbiota composition and molecular and metabolic resistance mechanisms on parasite infection dynamics is crucial for the design of novel transmission-blocking strategies.

No significant difference in the microbiota of deltamethrin exposed and unexposed mosquitoes was observed at 60 minutes or 72 hours post-exposure. One hour is likely insufficient time for the microbial composition to significantly shift in response to insecticide, given the relatively slow rate of bacterial growth, and the fact that any bacteria killed by insecticide exposure would still have been present during microbiota extraction. At 72 hours, differences in the microbiota of exposed and unexposed mosquitoes, if present, should have been apparent. The lack of difference observed may in part be due to the low sample size of the 5 day old 10x resistant group and may also reflect the short insecticide contact time. Bioassays are a single exposure at a lethal dose in a sterile environment used to determine resistance phenotype[51]. In the wild, multiple insecticide exposures are likely to happen at sub-lethal doses at both the larval stage, as habitats are contaminated with agricultural pesticides, and adult stage, as there is frequent interaction with treated surfaces or materials indoors[94]. Bioassays may therefore not induce the same shifts in microbiota as insecticide exposure in the wild. Furthermore, as mosquitoes acquire most of their microbiota from the aquatic environment at the larval stage[21], and may obtain insecticide metabolising bacteria at this stage, studying the effects of deltamethrin exposure on larvae, or adults which were exposed as larvae, may be more informative.

Our results demonstrated a significant relative reduction in alpha diversity in resistant and control 5-6 day old mosquitoes compared with the 2-3 day old group, as expected based on prior reports that microbial diversity declines with age[50]. Older mosquitoes also had lower relative abundances of *Ochrobactrum, Lysinibacillus* and *Stenotrophomonas*, the insecticide degrading species shown to be significantly enriched in resistant 2-3 day old individuals. *Serratia* and *Asaia,* the species associated with susceptibility, were present in increased abundance in the older age group. It has been widely reported that insecticide resistance declines with age[95]–[100]. The shift in microbiota may also be a contributing factor, and further research is warranted to determine the association between the microbiota, resistance phenotype and age.

In our study, qPCR detection of *Serratia* and *Asaia* was consistent with the 16S rRNA sequencing data, with susceptible mosquitoes having significantly lower CT values than resistant or control individuals. Population-level field screening using qPCR, as a cheaper, faster and more feasible option that amplicon sequencing, should be considered for integration in wider insecticide resistance monitoring, if reliable and reproducible bacterial markers associated with phenotype can be identified.

## Conclusions

Insecticide susceptibility is influenced by a range of diverse factors including host genetics, detoxification systems and behaviour as well as the mosquito microbiota. We report significant differences in the microbiota of deltamethrin-resistant *An. coluzzii* and have identified several bacterial species which were associated with either resistance or susceptibility to the host and therefore may represent important markers of resistance phenotype. The role of bacteria in determining resistance phenotype is highly complex and specific to the host and bacterial species, and insecticide and likely involves multiple, parallel mechanisms, including direct degradation of insecticide, altered host immune system, and changes to the midgut. In addition, these interactions may have important implications for host species fitness, vector competence and pathogen development and transmission. Further investigation into the mechanisms of microbiota mediated susceptibility is necessary as this may provide opportunities for preventing or reducing insecticide resistance, which is crucial to maintain gains in malaria vector control.

## Supporting information

Description of Supplementary Files

## Declarations

### Ethics approval and consent to participate

The study protocol was reviewed and approved by the Comite National d’Ethique des Sciences de la Vie et de la Sante (#069-19/MSHP/CNESVS-kp) and the institutional review board (IRB) of the London School of Hygiene and Tropical Medicine (#16860); all study procedures were performed in accordance with relevant guidelines and regulations. Prior to study initiation, community consent was sought from village leaders and written, informed consent was obtained from the heads of all households selected for participation and from all fieldworkers who performed HLCs. Fieldworkers participating in HLCs were provided with doxycycline malaria prophylaxis for the duration of the study.

### Consent for publication

Not applicable

### Availability of data and material

Sequence data generated by this study is available at Sequence Read Archive (SRA) BioProject PRJNA702915 (accession numbers: SRR13743435 – SRR13743530). All other relevant data are available from the corresponding author upon reasonable request.

Codes can be accessed at the public repository Zenodo (http://zenodo.org) under https://doi.org/10.5281/zenodo.4548776

### Author Contributions

ND, LAM, BP, and TW designed the study. BP, EC, AM and CE conducted fieldwork and BP, MK and LAM undertook mosquito rearing, phenotyping and preparation of samples for sequencing. Laboratory supervision was provided by LAM, CLJ and TW. BP, ND, and LAM performed the data analysis; and BP and LAM drafted the manuscript. All authors read and approved the final manuscript.

## Acknowledgements

The authors express their sincere thanks to Fidele Assamoa, Claver Adjobi and Laurent Didier Dobri, laboratory technicians at Centre Suisse de Recherches Scientifiques en Côte d’Ivoire (CSRS), for their support in mosquito collection and rearing; the chief and population of the village of Aboudé (Agboville), and the entomology fieldworkers of CSRS.

## Supplementary Information

Description of Supplementary Files

Table S1

Table S2

Table S3

Table S4

Table S5

Figure S1

Figure S2

Figure S3

Figure S4

