## Supplementary material for "Overabundance of *Asaia* and *Serratia* bacteria is associated with deltamethrin insecticide susceptibility in *Anopheles coluzzii* from Agboville, Côte d’Ivoire": Table S1

**Table S1:** Numbers of mosquitoes selected for DNA extraction and 16S rRNA sequencing. Mosquitoes were allocated to a concentration/time group based on the concentration of deltamethrin to which they were exposed, the period of follow up post exposure and whether they survived exposure. A selection from each group were pooled in groups of three and sequenced.

| Resistance phenotype | Description | Concentration  /time group | Age at death | Number of individuals | Number selected for sequencing (#pools) |
| --- | --- | --- | --- | --- | --- |
| Susceptible | Knocked-down at 60 minutes post exposure to the diagnostic dose (1x) deltamethrin. | Susceptible | 2-3 days | 97 | 87 (29) |
| Resistant | Survivors at 60 minutes post exposure to 5x deltamethrin | 5xRes60 | 2-3 days | 83 | 30 (10) |
|  | Survivors at 60 minutes post exposure to 10x deltamethrin | 10xRes60 | 2-3 days | 89 | 21(7) |
|  | Survivors at 72 hours post exposure to 5x deltamethrin | 5xRes72 | 5-6 days | 24 | 24(8) |
|  | Survivors at 72 hours post exposure to 10x deltamethrin | 10xRes72 | 5-6 days | 33 | 12 (4) |
|  | Survivors at 72 hours post exposure to diagnostic dose (1x) deltamethrin | 1xRes72 | 5-6 days | 55 | 36 (12) |
| Controls | Mosquitoes from control bottle which were alive 60 minutes after the initiation of bioassay | ControlA60 | 2-3 days | 57 | 42 (14) |
|  | Mosquitoes from control bottle which were alive 72 hours after the initiation of bioassay | ControlA72 | 5-6 days | 69 | 30 (10) |
| Blanks | 70% Ethanol which underwent same DNA extraction and 16S rRNA sequencing as samples | Ethanol blank | N/A | 1 | 1 (1) |
|  | Sterile, RNAse free water which underwent same DNA extraction and 16S rRNA sequencing as samples | Water blank |  | 3 | 3 (1) |
