## Supplementary figures and images for "Overabundance of *Asaia* and *Serratia* bacteria is associated with deltamethrin insecticide susceptibility in *Anopheles coluzzii* from Agboville, Côte d’Ivoire"

### Figure S1

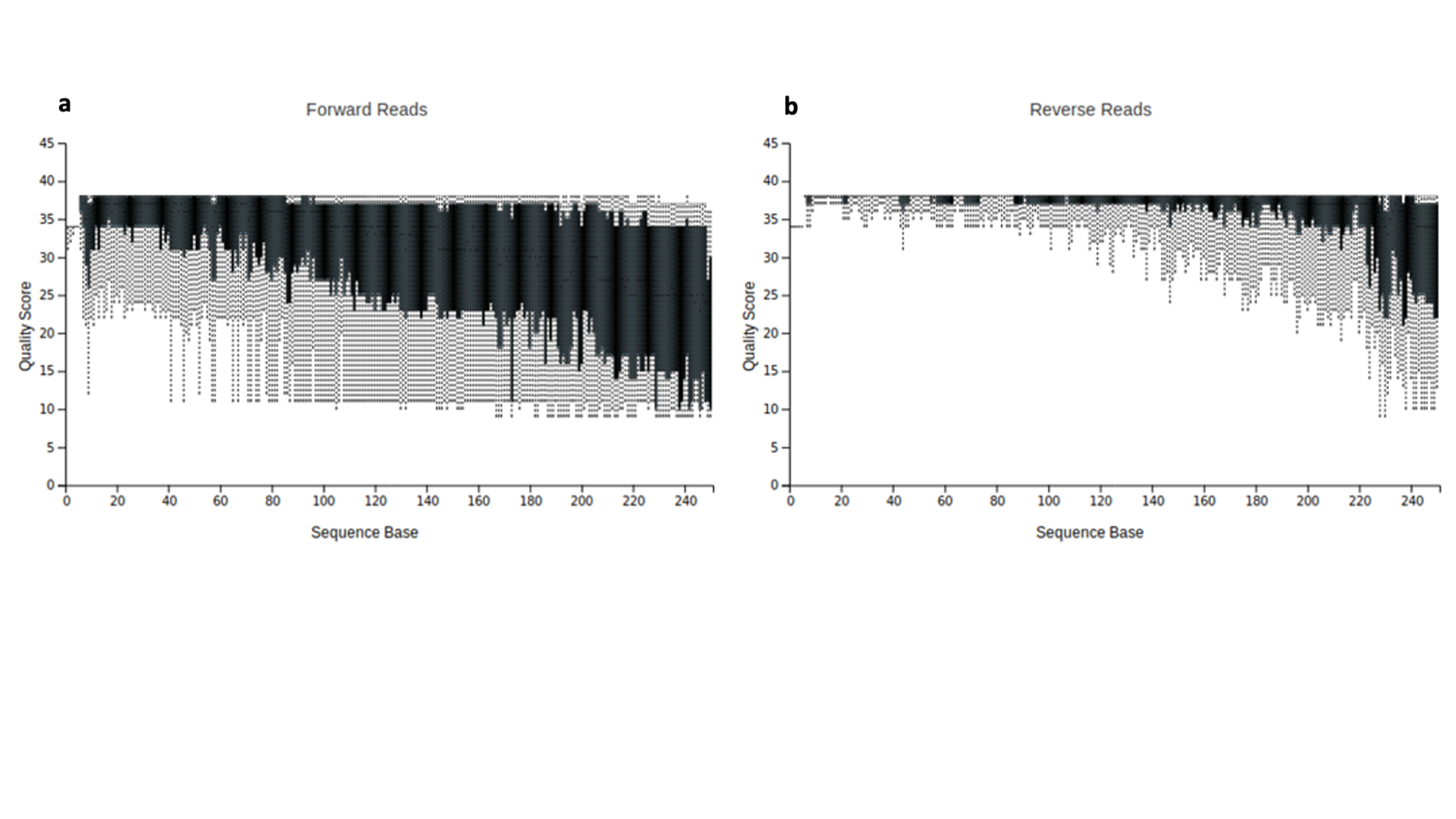

### Figure S2

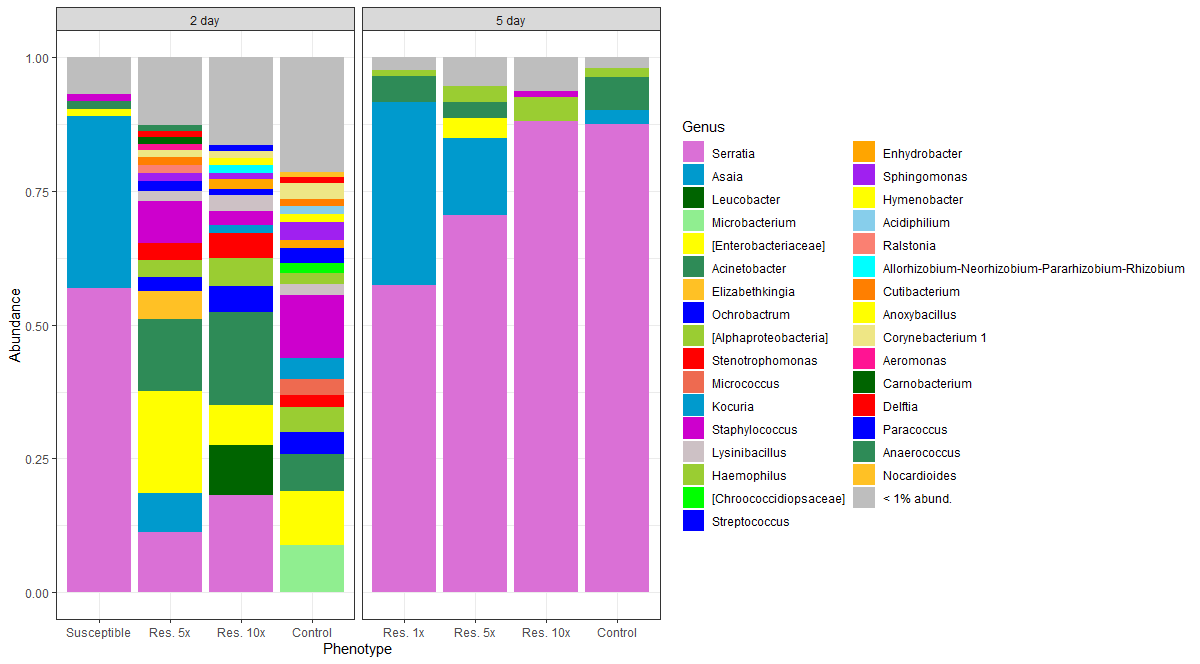

### Figure S3

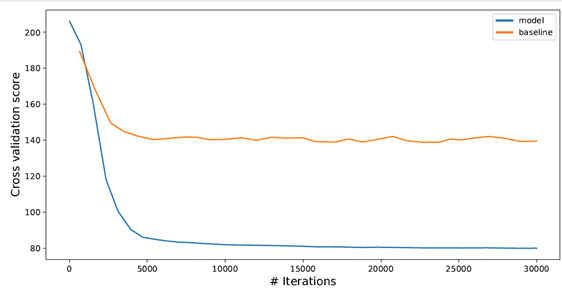

### Figure S4

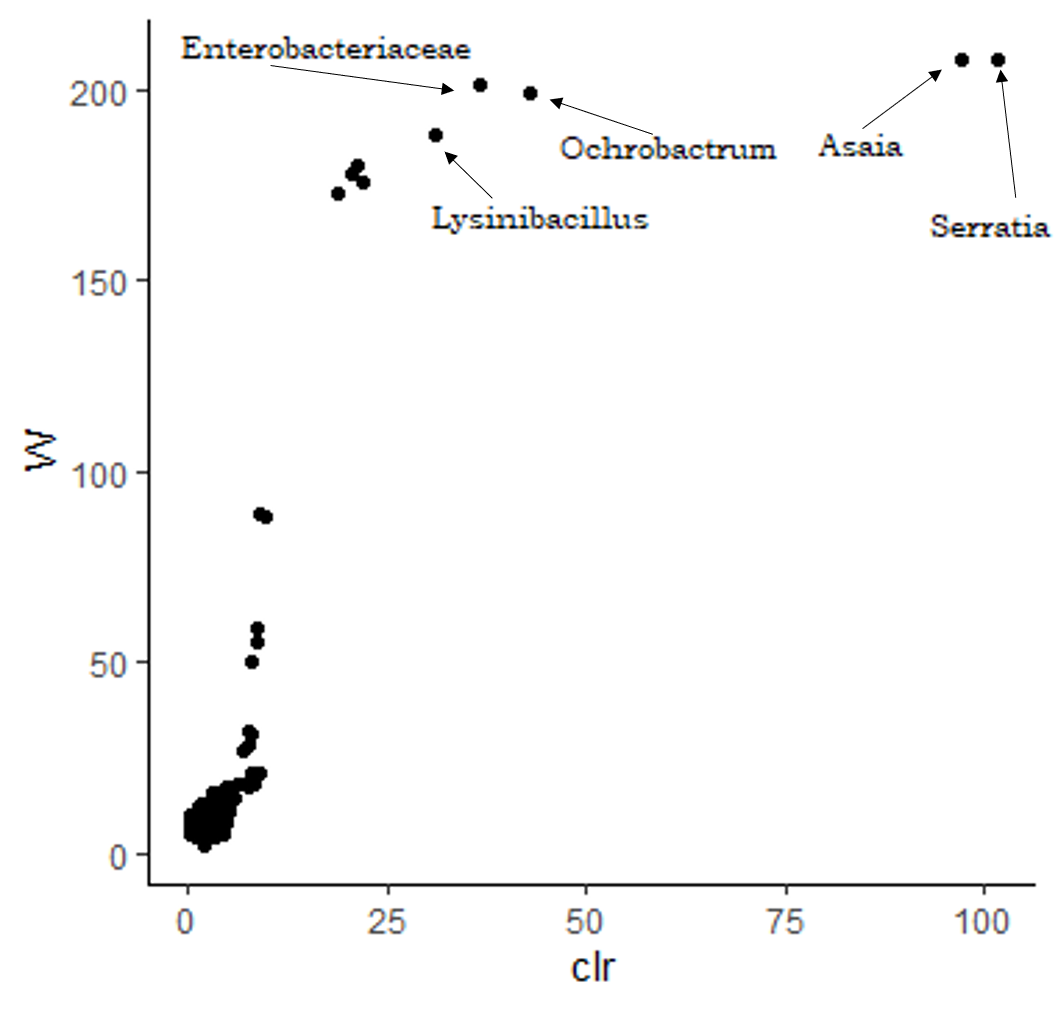
