## Supplementary material for "Overabundance of *Asaia* and *Serratia* bacteria is associated with deltamethrin insecticide susceptibility in *Anopheles coluzzii* from Agboville, Côte d’Ivoire": Description of Supplementary Files

File name: **Table S1.** **Numbers of mosquitoes selected for DNA extraction and 16S rRNA sequencing.** Mosquitoes were allocated to a concentration/time group based on the concentration of deltamethrin to which they were exposed, the period of follow up post exposure and whether they survived exposure. A selection from each group were pooled in groups of three and sequenced.

File name: **Table S2.** **Beta diversity outputs, showing no significant difference between rarefied and non-rarefied data.** Samples which fail to meet the rarefaction depth are excluded from analysis, therefore the sample sizes of rarefied data are reduced.

File name: **Table S3.** **Summary statistics of raw sequencing outputs and resulting data following denoising with Dada2 and filtering.**

File name: **Table S4. Relative abundances of bacterial species present in mosquitoes, grouped by age and resistance status.** Taxa are annotated to genus or lowest possible taxonomic level (square brackets).

File name: **Table S5. Differential rankings for features and taxa present in 2-3 day old *An. coluzzii,* as computed by Songbird.**

File name: **Figure S1. Quality plots for forward (a) and reverse (b) untrimmed reads, generated in Qiime2.** Low quality forward reads resulted in sequence loss due to failure of strands to merge during denoising, therefore only the reverse reads were used for downstream analysis.

File name: **Figure S2.** **Stacked bar plot showing relative abundance of taxa present in 2 day old and 5 day old *An. coluzzii* grouped by resistance phenotype.** Taxa are annotated to genus or lowest possible taxonomic level. Taxa present at less than 1% abundance are grouped together and coloured in grey.

File name: **Figure S3.** **Songbird model.** Blue line is trained on the dataset and is a predictor of model accuracy in contrast to the null hypothesis (orange). Illustrates the accuracy of model prediction and proportion of variance in microbial composition explained by resistance phenotype.

File name: **Figure S4.** **Volcano plot of differentially abundant bacterial taxa in 2-3 day old *An. coluzzii***, **as computed by analysis of composition of microbiomes (ANCOM).**
